# Selective Gut-Related Biomarker Profiles Define Parkinson Disease Trait and State

**DOI:** 10.64898/2026.11.12.766217

**Authors:** Ali Keshavarzian, Mohammad Samsul Alam, Phillip A. Engen, Robin M. Voigt, Maliha Shaikh, Ankur Naqib, Michelle Villanueva, Amanda Pires Bonfanti, Leila B. Giron, Neha Joshi, Stefan J. Green, Mohamed Abdel-Mohsen, Sheng Luo, Christopher G. Goetz

## Abstract

**Objective:** Gut-brain-axis abnormalities are increasingly implicated in Parkinson disease (PD), but most studies examine individual biological compartments of this axis. We asked whether readily accessible stool and plasma-based biomarker profiles show broad or selective gut abnormalities in PD and whether the profiles associated with PD diagnosis (Trait) are the same as those associated with clinical severity (State).

**Methods:** This cross-sectional study included 211 participants with PD and 204 healthy controls. Twenty-four stool and plasma biomarkers were organized into an overall profile, three parent domains (microbiome/dysbiosis, gut barrier/inflammation, systemic inflammation/immune activation), and eight more focused mechanistic profiles. Rank-based Global Statistical Tests (GSTs) compared PD Trait and State.

**Results:** The overall profile showed no evidence of a PD vs. control difference (GST P=0.607; permutation P=0.521), whereas the domain of microbiome/dysbiosis differed (GST P=0.017; permutation P=0.013). Four of eight focused profiles differed nominally: microbial functional-groups, microbial functional-ratios, epithelial injury/bacterial translocation, and myeloid activation. Across PD clinical severity, overall-profile results differed (GST P=0.053; permutation P=0.037); both categories of microbiome/dysbiosis (GST P=0.023; permutation P=0.016) and microbial alpha-diversity (GST P=0.027; permutation P=0.031) differed.

**Interpretation:** The analyses detected differences associated with PD Trait and State in selected gut-related profiles rather than a detectable overall PD-control difference. Microbiome-related profiles showed the most consistent domain findings across PD Trait and State, but they are sufficiently different to suggest that causation and progression of PD follow distinctly different gut processes. These findings refine the concept of the “Parkinson gut” and prioritize the microbiome for mechanistic and interventional studies.

## INTRODUCTION

A growing body of evidence links Parkinson disease (PD) to abnormalities across the gut-brain axis, including altered intestinal microbial communities and microbial metabolites, impaired intestinal barrier integrity, epithelial injury, and intestinal and systemic inflammation.^1–9^ These observations have generated the concept of a “Parkinson gut,” but they have not established whether gut-related abnormalities form a single coordinated phenotype or instead represent selective alterations affecting different biological processes.^2, 3, 5, 7, 10^

This distinction matters for biomarker development. Direct interrogation of intestinal tissue can provide biological depth, but biopsies are invasive and difficult to repeat at scale. Stool and blood measurements are substantially more accessible and can provide complementary readouts of microbial composition, epithelial injury, microbial-product exposure, mucosal inflammation, and systemic immune activation. The key translational question is therefore not which compartment is biologically “most important,” but whether accessible peripheral measurements capture a broad PD-associated signal or selective biomarker profiles that may be useful for future longitudinal studies and therapeutic target development.

A second question is whether gut-related profiles associated with PD diagnosis (Trait) are the same as those associated with clinical severity (State) among people who already have PD. A biomarker profile that distinguishes PD from controls may not necessarily worsen with advancing clinical state, and a state-sensitive profile may be different one that distinguishes PD from controls. Separating these mechanistic questions is important for defining future biomarker contexts of use, including studies of disease risk and onset, progression, patient stratification, and treatment response.

We addressed these questions in 211 participants with PD and 204 neurologically healthy controls using 24 unique stool and plasma biomarkers, considered as a single overall gut-related profile [Level 1 analysis], then divided and analyzed by three broad mechanistic groups [Level 2 analysis] spanning microbiome/dysbiosis, mucosa gut leakage/inflammation, and systemic inflammation/immune activation, and finally eight more focused mechanistic clusters [Level 3 analysis]. Secondary analyses explored ordered study-defined clinical state, Hoehn and Yahr (H&Y) stage, MDS-UPDRS Part III motor scores, and plasma neurofilament light chain (NfL) burden.

## METHODS

### Study Design and Study Participants

This was a mechanistically based, observational, cross-sectional analysis of prospectively collected repository samples and clinical data, approved by the Rush University Medical Center (RUMC) Institutional Review Board (IRB: ORA#23102401, ORA#23011304, ORA#10062805). All participants provided written informed consent, and all procedures were conducted in accordance with institutional guidelines and the Declaration of Helsinki. Clinical information was collected prospectively using structured and/or validated instruments. Laboratory personnel were blinded to disease group and study-defined PD clinical-state group, and the initial Rank-based Global Statistical Tests (GSTs) calculations were completed while group codes were masked. The study included 204 community controls and 211 participants with PD (**Fig 1A**; **Table 1**). A paired subgroup analysis included 48 of these PD participants and their household spousal controls (48 pairs; **Table S1**).

**Figure 1.**
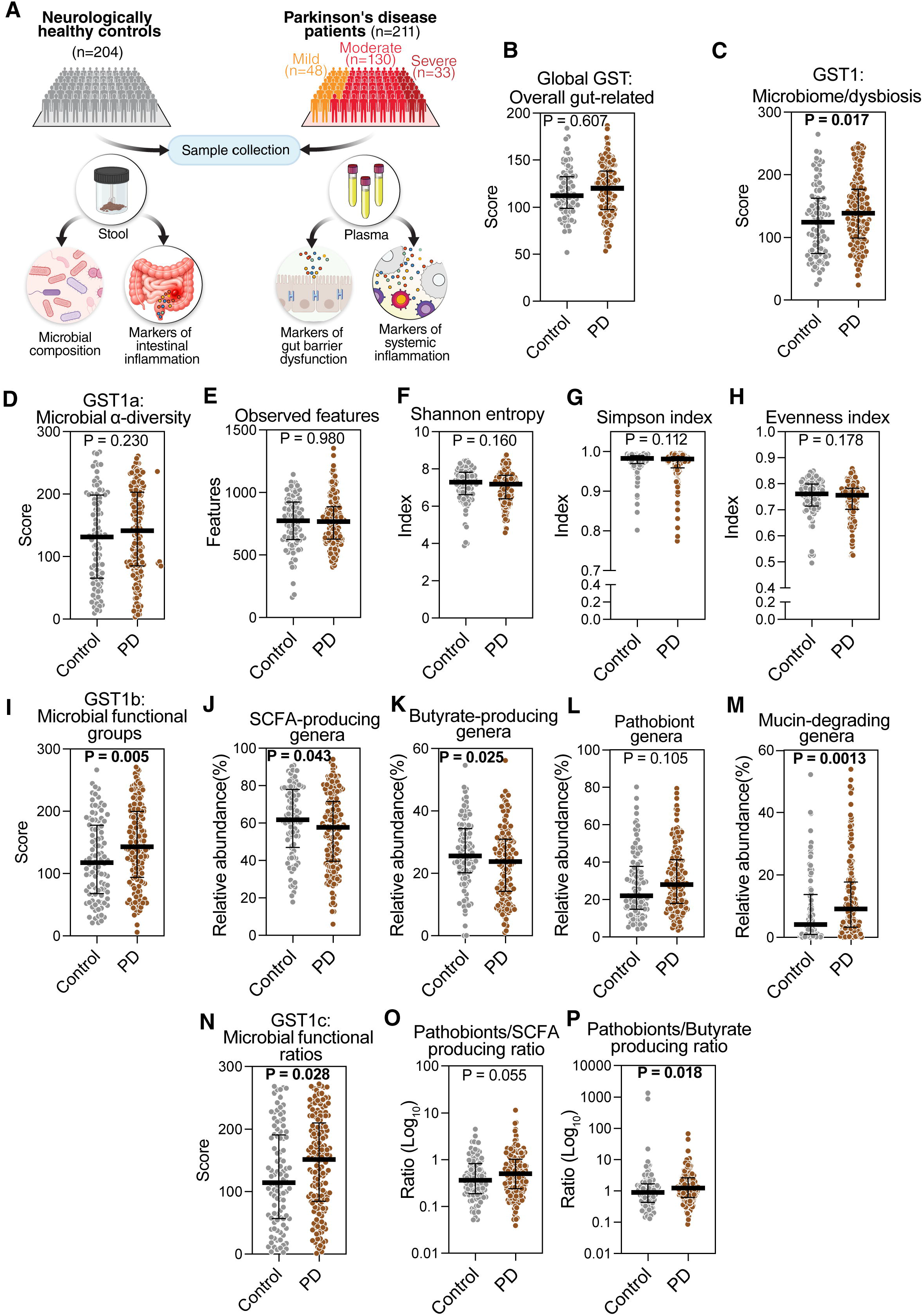
Study design and microbiome-related biomarker profiles according to PD status. (**A**) Study design and sample sources. (**B**) Overall gut-related 24-biomarker GST profile. (**C**) Broad mechanistic microbiome/dysbiosis profile. (**D-H**) Focused microbial alpha-diversity profile and constituent biomarkers. (**I-M**) Focused microbial functional-group profile and constituent biomarkers. (**N-P**) Derived microbial functional-ratio profile and constituent ratios. Higher GST scores indicate greater prespecified PD-associated abnormality after directional orientation and rank transformation. The functional-ratios were not included in Level 1 or Level 2 because they are derived from component functional-group measures. (**B–D**, **I**, **N**) P-values are from parametric GSTs for profile panels; (**E–H**, **J–M**, **O–P**) P-values are from Mann–Whitney U tests for constituent biomarkers. Bars show medians with interquartile ranges. Sample sizes (control, PD): (**B**) 86, 147; (**C–H**) 97, 173; (**I–P**) 100, 173.

**Table 1.** Demographic and clinical characteristics of study participants overall and according to PD status and clinical state.

| Characteristics | Overall<br>N=415 | Control<br>N=204 | PD |  |  | All PD<br>N=211 |
|---|---|---|---|---|---|---|
|  |  |  | Mild<br>N=48 | Moderate<br>N=130 | Severe<br>N=33 |  |
| Age, years | 61 [15] | 57.5 [13] | 64 [13] | 70 [15] | 65 [13] | 65 [11.5] |
| Sex, n (%) |  |  |  |  |  |  |
| Male | 231 (55.7) | 92 (45.1) | 31 (64.6) | 87 (66.9) | 21 (63.6) | 139 (65.9) |
| Female | 184 (44.3) | 112 (54.9) | 17 (35.4) | 43 (33.1) | 12 (36.4) | 72 (34.1) |
| Race, n (%) |  |  |  |  |  |  |
| White | 331 (79.8) | 132 (64.7) | 44 (91.7) | 127 (97.7) | 28 (84.8) | 199 (94.3) |
| Black | 62 (14.9) | 57 (27.9) | 0 (0.0) | 2 (1.5) | 3 (9.1) | 5 (2.4) |
| Asian | 12 (2.9) | 7 (3.4) | 3 (6.2) | 1 (0.8) | 1 (3.0) | 5 (2.4) |
| Other | 10 (2.4) | 8 (3.9) | 1 (2.1) | 0 (0.0) | 1 (3.0) | 2 (0.9) |
| Ethnicity, n (%) |  |  |  |  |  |  |
| Non-Hispanic or Latino | 392 (94.5) | 192 (94.1) | 47 (97.9) | 123 (94.6) | 30 (90.9) | 200 (94.8) |
| Hispanic or Latino | 23 (5.5) | 12 (5.9) | 1 (2.1) | 7 (5.4) | 3 (9.1) | 11 (5.2) |
| Clinical |  |  |  |  |  |  |
| Body Mass Index (BMI) | 26.58 [6.60] | 28.32 [6.37] | 25.70 [5.30] | 26.50 [5.55] | 24.73 [8.40] | 26.12 [5.90] |
| Bristol Stool Scale Score | 3.50 [2.00] | 4.00 [1.00] | 4.00 [3.00] | 2.00 [2.00] | 1.00 [1.00] | 2.00 [3.00] |
| Disease Duration (years) | - | - | 2.00 [3.75] | 6.50 [6.50] | 10.00 [7.50] | 6.00 [7.00] |
| Hoehn & Yahr Staging | - | - | 2.00 [0.00] | 2.00 [0.00] | 3.00 [0.00] | 2.00 [0.00] |
| MDS-UPDRS: Part 3 | - | - | 23.50 [16.50] | 25.00 [18.25] | 36.00 [22.25] | 25.00 [18.00] |
| Levodopa Daily Dosage (mg/day) | - | - | - | 400 [300] | 500 [475] | 300 [550] |

Abbreviations: H&Y, Hoehn and Yahr; IQR, interquartile range; PD, Parkinson disease; MDS-UPDRS, Movement Disorder Society–Unified Parkinson’s Disease Rating Scale. Continuous variables are summarized as median [IQR], and categorical variables as n (%), with percentages calculated using group-specific denominators. Mild PD was defined as Hoehn and Yahr stage 1-2 and PD medication-naive; moderate PD as Hoehn and Yahr stage 1-2 while receiving a stable dose of PD medication; and severe PD as Hoehn and Yahr stage 3-5 while receiving a stable dose of PD medication. Because this classification incorporates both H&Y stage and medication status, these groups represent study-defined clinical states rather than pure disease-severity strata.

Participants with PD, defined by Movement Disorder Society criteria,^11^ and were classified into three study-defined clinical states: 48 mild (H&Y stages 1-2 and medication-naive), 130 moderate (H&Y stages 1-2 but receiving a stable dose of PD medication), and 33 severe (H&Y stages 3-5 while receiving a stable dose of PD medication).^12^ Because this classification incorporates both H&Y stage and medication status, it was treated as a study-defined clinical-state construct rather than a pure disease-severity scale. H&Y stage and MDS-UPDRS Part III Motor Examination scores were assessed on the day of sample collection; H&Y and MDS-UPDRS Part III were obtained in the ON-medication state where applicable.

### Specimen Collection and Microbiome Profiling

Detailed protocols for blood (plasma) and stool collection, specimen processing and storage, and stool microbiome profiling, including DNA extraction, near-full-length 16S rRNA gene sequencing, and bioinformatic processing, are provided in the **Supplementary Methods**.

### Gut Microbiome and Dysbiosis Biomarkers

The broad mechanistic gut microbiome/dysbiosis domain encompassed eight biomarkers: four microbial alpha-diversity metrics and four microbial functional-groups. Alpha-diversity metrics included Shannon and Simpson indices, observed features, and Pielou’s evenness.^13^ For the microbial functional-groups, bacterial genera were classified as putative short-chain fatty acid (SCFA)-producing, butyrate-producing, pathobiont-associated, or mucin-degrading taxa based on published literature.^14–16^ Genus-level relative abundances, obtained from the taxonomic BIOM file, were added within each functional-group for each participant’s fecal sample. Additionally, a focused mechanistic subset of microbial functional-ratios was examined, calculated by dividing the total relative abundance of putative pathobiont-associated genera by that of SCFA-producing or butyrate-producing genera. The bacterial genera included in these analytic profiles are summarized in the **Source Data**.

### Mucosa Gut Leakage and Inflammatory Biomarkers

The broad mechanistic mucosa gut leakage and inflammation profile comprised six biomarkers. Three plasma biomarkers for mucosa gut leakage (intestinal epithelial injury and bacterial translocation) were assessed: lipopolysaccharide-binding protein (LBP),^3, 17^ intestinal fatty acid-binding protein (I-FABP),^18, 19^ and regenerating islet-derived protein 3 alpha (REG3A).^20, 21^ Concentrations were measured using the Human DuoSet ELISA Development System for LBP (ng/mL) (R&D Systems/Bio-Techne, #DY870-05), Human FABP2/I-FABP (pg/mL) Quantikine ELISA Kit (R&D Systems/Bio-Techne, #DFBP20), and Human REG3A (pg/mL) ELISA Kit (RayBiotech, #ELH-REG3A-1). Two plasma and one stool based biomarker of gut associated inflammation were assessed: interleukin-8 (IL-8/CXCL8) and interleukin-17A (IL-17A) as circulating inflammatory markers of gut inflammation,^22, 23^ and stool calprotectin.^7, 24^ Concentrations were measured for IL-8 (pg/mL) using U-PLEX Custom Immuno-Oncology Group 1 Meso Scale Discovery (MSD, #K151AEM-2), IL-17A (pg/mL) using U-PLEX Custom Metabolic Group 1 (MSD, #K151ACM-2), and stool calprotectin (µg/g) using the BÜHLMANN assay (#EK-CAL2). All assays were performed according to the manufacturers’ instructions. Additionally, plasma zonulin concentrations (ng/mL) were measured using the Human Zonulin ELISA Kit (MyBioSource, MBS167049).

### Systemic Inflammation and Immune Activation Biomarkers

The broad mechanistic systemic inflammation and immune activation domain included ten plasma biomarkers that were measured using electrochemiluminescence-based multiplex immunoassays and ELISA. Markers of myeloid activation included soluble CD14 (sCD14; R&D Systems/Bio-Techne, #DC140), soluble CD163 (sCD163; R&D Systems/Bio-Techne, #DY1607), and soluble urokinase plasminogen activator receptor (uPAR; MSD, #K151AEM-2).^25–27^ Markers of innate cytokine inflammation included interleukin-1 beta (IL-1β) and interleukin-6 (IL-6), both measured using U-PLEX Custom Metabolic Group 1 (MSD, #K151ACM-2), and tumor necrosis factor-alpha (TNF-α; MSD, #K151AEM-2).^28, 29^ Markers of the interferon-gamma (IFN-γ)-associated inflammatory signaling included interleukin-12p70 (IL-12p70; MSD, #K151ACM-2), IFN-γ, CXCL9, and CXCL10 (MSD, #K151AEM-2).^28, 30–33^.

All assays followed the manufacturers’ instructions, and plasma concentrations were reported in pg/mL. Values below the limit of detection ( LOD), which occurred only for IL-1β, TNF-α, IL-12p70, and IFN-γ, were quality controlled assigned 50% of the lowest measured value for the corresponding biomarker, and colored coded in their respective figures.

### Plasma Biomarker of Axonal Neurodegeneration

Plasma neurofilament light chain (NfL) concentrations (pg/mL), a biomarker of axonal neurodegeneration and injury,^34^ were measured using the R-PLEX Human Neurofilament L Assay (MSD, K1517XR-2) according to the manufacturer’s instructions.

### Statistical Analysis

Rank-based Global Statistical Tests (GSTs) were used to compare PD vs healthy controls (PD Trait) and to compare different PD severity categories (PD State). Twenty-four unique stool and plasma biomarkers were organized into three levels of GST analysis: an overall gut-related profile [Level 1 GST analysis; 24 biomarkers]; three non-overlapping broad mechanistic domains [Level 2 GST analysis] including microbiome/dysbiosis (8 biomarkers), mucosa gut leakage/inflammation (6 biomarkers), and systemic inflammation/immune activation (10 biomarkers); and eight more focused biologically mechanistic clusters [Level 3 GST analysis] (**Table S2**). Derived microbial functional-ratios were excluded from Levels 1 and 2 GST analyses because the component quantities were already represented. Analytic samples varied by available biomarkers; these levels were not interpreted as nested tests based on one common complete-case sample.

The rank-based GST framework followed O’Brien’s approach,^35^ and its previous use in PD.^36^ Biomarkers were oriented so higher values represented greater hypothesized PD-associated abnormalities; favorable-direction variables were reverse-oriented. Values below the detection limit were retained as tied observations below the lowest measurable value and assigned their average lower rank. Participants with complete constituent-marker data and an observed group assignment were included for each GST profile, with ranks recalculated within that profile’s analytic sample. The GST score was the mean of its constituent marker ranks. For secondary regression analyses, scores were first computed among participants with complete constituent-marker data; regression-specific complete-case exclusions were applied subsequently.

Two-group GST comparisons were based on two-sample *t*-tests expressed as F=t²; comparisons across three clinical-state groups used one-way analysis of variance. Kruskal– Wallis and permutation analyses were robustness checks, with 10,000 label permutations for unpaired comparisons. For paired PD–spouse analyses, permutation tests randomly reversed the sign of each within-pair difference, using a two-sided mean-difference statistic and plus-one correction. GST group tests were unadjusted for demographic covariates. All P-values were two-sided and unadjusted for multiple comparisons; Analysis for eight more focused mechanistic clusters [Level 3 GST analysis], individual-marker, and sensitivity results were exploratory. Neither a nonsignificant broader test nor a nominally significant focused test was interpreted as proof that the corresponding biological domain was normal or abnormal.

Within PD, exploratory secondary analyses treated study-defined clinical state as an ordered variable (mild=0, moderate=1, severe=2). Separate age- and sex-adjusted linear models used each standardized focused profile as the outcome, and a complete-case multivariate response model jointly included all eight focused profiles (**Tables S3-S4**). A spouse-paired sensitivity analysis used paired t-tests, Wilcoxon signed-rank tests, and two-sided sign-flip permutation tests in available PD-spouse dyads (**Tables S5-6**).

Alternative exploratory clinical characterizations included: (1) GST comparisons by H&Y stage 1–2 versus 3–5 (**Tables S7-8**); (2) age- and sex-adjusted regressions of continuous MDS-UPDRS Part III on standardized gut profiles, with D3 excluded from the joint focused-profile model because it is constructed from quantities used in D2 (**Table S9**); and (3) unadjusted GST comparisons among low, intermediate, and high cohort-defined NfL groups (**Tables S10-11**). The NfL analyses were cross-sectional associations, not longitudinal assessments of disease progression. All analyses were conducted in R version 4.6.0.

We excluded zonulin values from the final GST analysis following the initial blinded GST analyses and assay quality-control review. We identified approximately two-fold higher zonulin concentrations on ELISA plates 2 and 6 compared to the other plates. These inter-plate differences, together with published concerns regarding the molecular specificity of commercial zonulin assays and their reliability as measures of intestinal permeability,^37–41^ prompted the exclusion of Zonulin from the final GST analysis. However, zonulin results (100 control, 126 PD-29 mild, 75 moderate, 22 severe) are reported separately as an exploratory biomarker (**Fig S1**).

## RESULTS

### Participant Characteristics

The study included 415 participants: 204 neurologically healthy controls and 211 participants with PD, comprising 48 mild, 130 moderate, and 33 severe study-defined clinical states (**Fig 1A**; **Table 1**). Compared with controls, the PD group was older (median 65 versus 57.5 years), included a higher proportion of men (65.9% versus 45.1%), and had a different racial distribution (94.3% versus 64.7% White; 2.4% versus 27.9% Black or African American). Stool consistency also differed descriptively: the median Bristol Stool Scale score was 2 in PD versus 4 in controls and declined from 4 in mild to 2 in moderate and 1 in severe PD. Across the study-defined PD states, median disease duration increased from 2.0 to 6.5 to 10.0 years and median MDS-UPDRS Part III increased from 23.5 to 25.0 to 36.0, supporting increasing clinical burden across the PD groups while underscoring that the construct also incorporates medication status.

### PD Diagnosis (Trait): Overall and Microbiome Profiles

The central PD-control finding was selectivity of gut involvement rather than a uniform gut-related shift. The overall 24-biomarker GST gut-related profile did not differ between PD and controls (GST P=0.607; permutation P=0.521; **Fig 1B**; **Table 2**). In contrast, the broad mechanistic domain of microbiome/dysbiosis differed (GST P=0.017; permutation P=0.013; **Fig 1C**; **Table 2**). This broad microbiome signal was not explained by alpha-diversity, whose focused profile was not different (GST P=0.230; **Fig 1D**; **Table 2**), and none of its four constituent biomarkers differed significantly between PD and controls (observed features P=0.980, Shannon index P=0.160, Simpson index P=0.112, evenness P=0.178; **Fig 1E-H**). Instead, the PD-control microbiome signal was concentrated in functional composition. The microbial functional-groups profile differed between PD and controls (GST P=0.005; permutation P=0.004; **Fig 1I**; **Table 2**). PD was characterized by lower total relative abundance of putative anti-inflammatory, SCFA-producing genera (P=0.043) and butyrate-producing genera (P=0.025), together with higher relative abundance of putative total mucin-degrading genera (P=0.0013); the putative total pathobiont genera did not differ (P=0.105) (**Fig 1J-M**). Consistent with this pattern, the derived microbial functional-ratio profile also differed (GST P=0.028; permutation P=0.027; **Fig 1N**; **Table 2**), with a higher pathobiont-to-butyrate-producing ratio in PD (P=0.018) and a borderline difference in the pathobiont-to-SCFA-producing ratio (P=0.055) (**Fig 1O-P**).

**Table 2.** GST results for Level 1 overall gut-related, Level 2 broad mechanistic domain, and Level 3 focused mechanistic profiles by PD trait and state.

| Level | Biological domain | Markers | GST F | GST P | KW H | Permutation P | Analytical n |
|---|---|---|---|---|---|---|---|
| **A. PD Trait (PD vs Control)** |  |  |  |  |  |  |  |
| 1 | Overall gut-related | 24 | 0.26 | 0.607 | 0.42 | 0.521 | 233 (147,86) |
| 2 | Microbiome/dysbiosis | 8 | 5.76 | **0.017** | 5.93 | **0.013** | 270 (173,97) |
| 3 | Microbial alpha-diversity | 4 | 1.44 | 0.230 | 1.67 | 0.200 | 270 (173,97) |
|  | Microbial functional groups | 4 | 7.95 | **0.005** | 7.65 | **0.004** | 273 (173,100) |
|  | Microbial functional ratios | 2 | 4.87 | **0.028** | 4.75 | **0.027** | 273 (173,100) |
| 2 | Mucosa gut leakage/inflammation | 6 | 0.05 | 0.832 | 0.11 | 0.736 | 235 (147,88) |
| 3 | Epithelial injury and bacterial translocation | 3 | 11.58 | **<0.001** | 11.07 | **<0.001** | 350 (175,175) |
|  | Intestinal mucosal inflammation | 3 | 0.00 | 0.986 | 0.01 | 0.912 | 235 (147,88) |
| 2 | Systemic inflammation/immune activation | 10 | 0.17 | 0.678 | 0.04 | 0.851 | 280 (163,117) |
| 3 | Myeloid activation | 3 | 9.02 | **0.003** | 8.23 | **0.004** | 350 (175,175) |
|  | Innate cytokine inflammation | 3 | 0.07 | 0.789 | 0.02 | 0.891 | 282 (165,117) |
|  | IFN-γ/CXCR3-axis inflammation | 4 | 0.38 | 0.541 | 0.63 | 0.430 | 280 (163,117) |
| **B. PD State (Mild, Moderate, Severe)** |  |  |  |  |  |  |  |
| **Level** | **Biological domain** | **Markers** | **GST F** | **GST P** | **KW H** | **Permutation P** | **Analytical n** |
| 1 | Overall gut-related | 24 | 2.99 | 0.053 | 6.64 | **0.037** | 147 (39,86,22) |
| 2 | Microbiome/dysbiosis | 8 | 3.88 | **0.023** | 8.08 | **0.016** | 173 (43,105,25) |
| 3 | Microbial alpha-diversity | 4 | 3.69 | **0.027** | 6.92 | **0.031** | 173 (43,105,25) |
|  | Microbial functional groups | 4 | 2.26 | 0.108 | 4.47 | 0.105 | 173 (43,105,25) |
|  | Microbial functional ratios | 2 | 1.98 | 0.141 | 3.63 | 0.162 | 173 (43,105,25) |
| 2 | Mucosa gut leakage/inflammation | 6 | 2.63 | 0.076 | 4.80 | 0.093 | 147 (39,86,22) |
| 3 | Epithelial injury and bacterial translocation | 3 | 1.85 | 0.161 | 2.77 | 0.253 | 175 (44,105,26) |
|  | Intestinal mucosal inflammation | 3 | 1.57 | 0.212 | 2.73 | 0.264 | 147 (39,86,22) |
| 2 | Systemic inflammation/immune activation | 10 | 1.50 | 0.226 | 2.74 | 0.261 | 163 (40,97,26) |
| 3 | Myeloid activation | 3 | 0.56 | 0.571 | 1.00 | 0.612 | 175 (44,105,26) |
|  | Innate cytokine inflammation | 3 | 1.10 | 0.334 | 3.00 | 0.226 | 165 (41,98,26) |
|  | IFN-γ/CXCR3-axis inflammation | 4 | 1.48 | 0.230 | 2.93 | 0.230 | 163 (40,97,26) |

Abbreviations: GST, global statistical test; KW, Kruskal–Wallis; PD, Parkinson disease.
Level 1 comprised an overall gut-related profile of 24 unique biomarkers. Level 2 partitioned these biomarkers into three nonoverlapping broad biological domains: microbiome/dysbiosis (8 biomarkers), mucosa gut leakage/inflammation (6 biomarkers), and systemic inflammation/immune activation (10 biomarkers). The eight Level 3 focused profiles are defined in Supplementary Table S2. The two derived microbial functional ratios were included only at Level 3 and excluded from Levels 1 and 2 to avoid double-counting their component measures. For PD-control and PD clinical-state comparisons, the GST F statistic is the parametric group-comparison statistic, and Kruskal–Wallis H is the nonparametric statistic. Permutation P values are reported as a robustness check. All P values are two-sided and unadjusted for multiple comparisons. GST group comparisons were unadjusted for demographic covariates. Analytic sample sizes vary across profiles because of component-biomarker availability; n is shown as total (PD, control) for PD-control comparisons and total (mild, moderate, severe) for clinical-state comparisons. Biomarkers were equally weighted after orientation and rank transformation. Level 3 focused-profile analyses are exploratory. For the four focused profiles with permutation P<0.05 in PD-control comparisons, oriented profile scores were higher in PD than controls, consistent with the prespecified adverse direction.

### PD Diagnosis (Trait): Mucosa Gut Leakage/Inflammation

The broad mucosa gut leakage/inflammation domain did not differ between PD and controls (GST P=0.832; permutation P=0.736; **Fig 2A**; **Table 2**), but its two focused clusters behaved differently: The intestinal epithelial injury/bacterial-translocation profile was higher in PD (GST P<0.001; permutation P<0.001; **Fig 2B**; **Table 2**), driven most clearly by higher plasma REG3A (P=0.0009), whereas I-FABP (P=0.091) and LBP (P=0.098) did not individually reach P<0.05 (**Fig 2C-E**). By contrast, the intestinal mucosal-inflammation profile did not differ (GST P=0.986; **Fig 2F**; **Table 2**). Within that profile, constituent biomarker stool calprotectin was higher in PD (P=0.015), while plasma IL-8 (P=0.953) and IL-17A (P=0.826) were not different (**Fig 2G-I**). Thus, focused epithelial-injury and stool-inflammatory signals were present despite the absence of a coordinated shift across the broader mucosa gut leakage/inflammation domain. However, the focused differences should not be interpreted as formal decompositions of the broad parent domains GST analysis result. Concerning the exploratory plasma zonulin analysis, separate zonulin comparisons did not reach nominal P<0.05 for PD versus controls (P=0.061) or across three study-defined PD clinical-state groups (P=0.774; **Fig S1A-B**).

**Figure 2.**
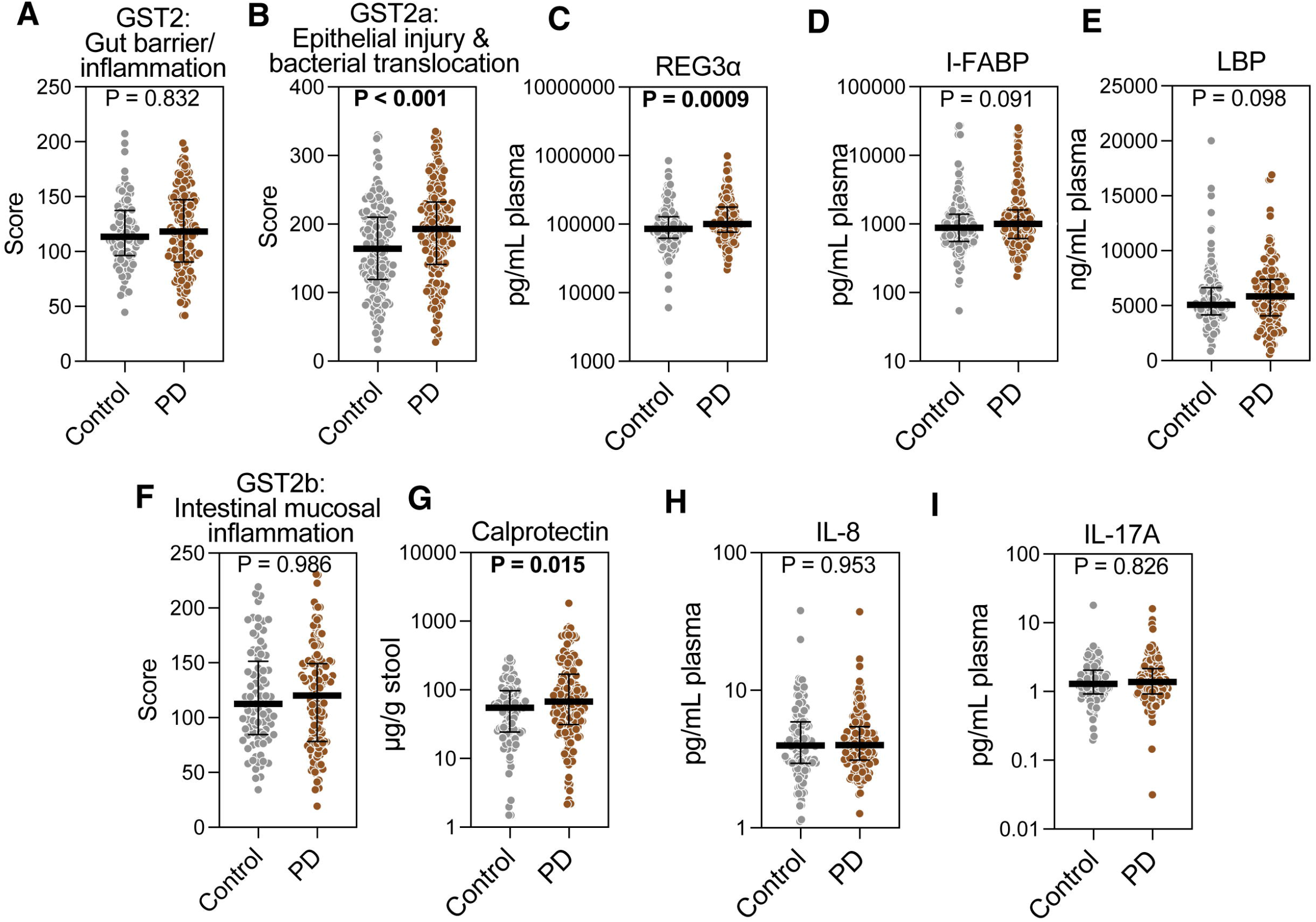
Mucosa gut leakage and inflammatory biomarker profiles according to PD status. (**A**) Broad mechanistic mucosa gut leakage/inflammation profile. (**B-E**) Focused epithelial injury/bacterial-translocation profile and constituent biomarkers of plasma REG3A, I-FABP, and LBP. (**F-I**) Focused intestinal mucosal inflammation profile and constituent biomarkers of stool calprotectin, plasma IL-8, and plasma IL-17A. Higher GST scores indicate greater prespecified PD-associated abnormality. (**A-B, F**) P-values are from parametric GSTs for profile panels; (**C-E**, **G-I**) P-values are from Mann–Whitney U tests for constituent biomarkers. Bars show medians with interquartile ranges. Sample sizes (control, PD): (**A, F**) 88, 147; (**B–E**) 175, 175; (**G**) 100, 173; (**H-I)** 117, 163.

### PD Diagnosis (Trait): Systemic Inflammation/Immune Activation

A similar nested pattern was observed for systemic inflammation. The broad systemic inflammation/immune-activation domain did not differ between PD and controls (GST P=0.678; permutation P=0.851; **Fig 3A**; **Table 2**), but the focused myeloid-activation profile was higher in PD (GST P=0.003; permutation P=0.004; **Fig 3B**; **Table 2**). Among its constituent biomarkers, sCD163 was clearly higher in PD (P<0.0001), uPAR showed a borderline difference (P=0.058), and sCD14 did not differ (P=0.938) (**Fig 3C-E**). Neither the innate-cytokine inflammation profile (GST P=0.789; **Fig 3F**) nor the IFN-γ/CXCR3-axis inflammation profile (GST P=0.541; **Fig 3J**, **Table 2**) differed, and no constituent cytokines in these two profiles reached P<0.05 (**Fig 3G-I, K-N**). Thus, selected focused profiles differed even though the corresponding the broad parent domains GST analysis did not.

**Figure 3.**
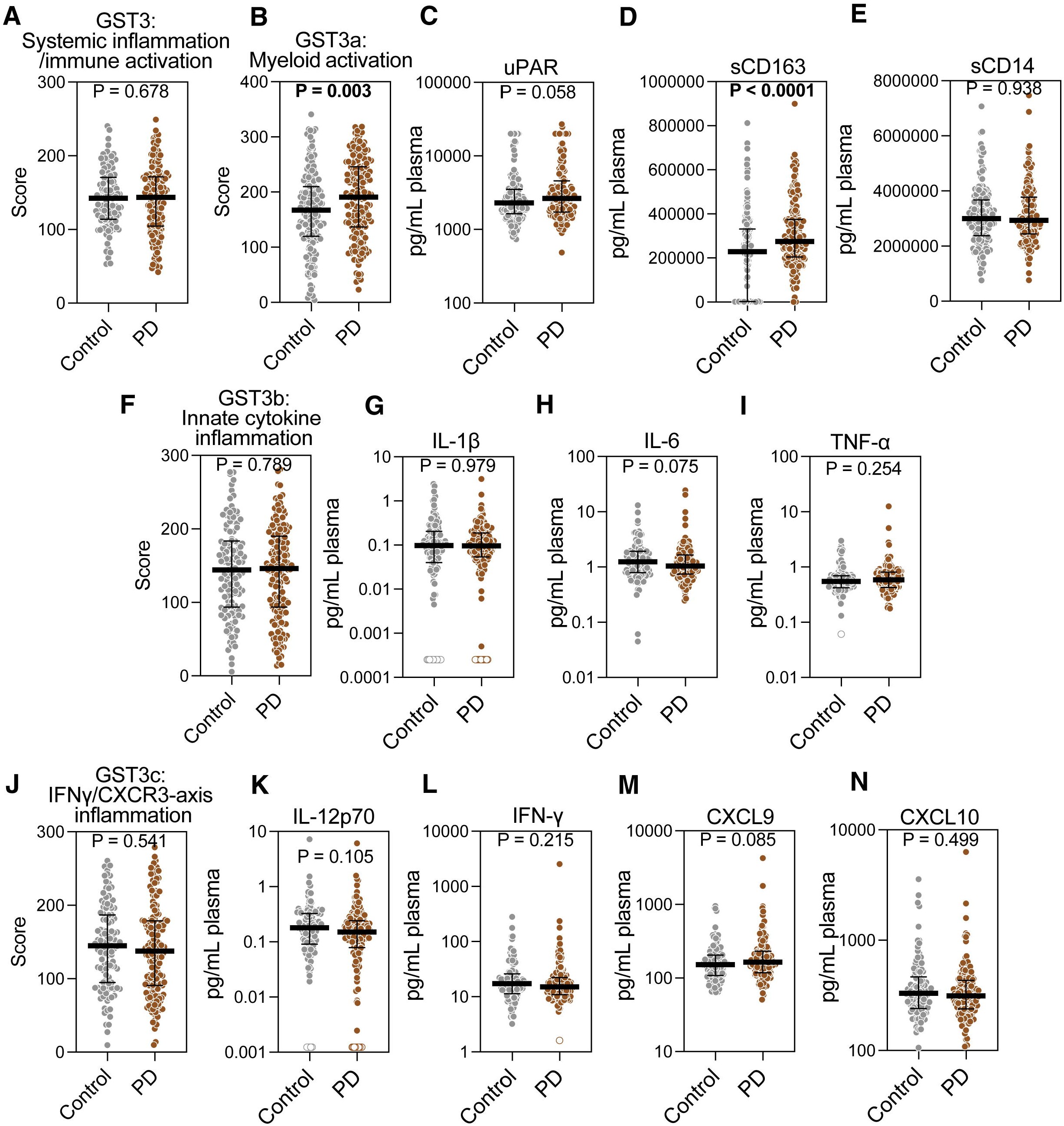
Systemic inflammatory and immune-activation biomarker profiles according to PD status. (**A**) Broad mechanistic systemic inflammation/immune-activation profile. (**B-E**) Focused myeloid-activation profile and constituent biomarkers. (**F-I**) Focused innate-cytokine inflammation profile and constituent biomarkers. (**J-N**) Focused IFN-γ/CXCR3-axis inflammation profile and constituent biomarkers. Higher GST scores indicate greater prespecified PD-associated abnormality. Values below the limit of detection ( LOD), for IL-1β, TNF-α, IL-12p70, and IFN-γ are colored coded in their respective figures. (**A-B, F, J**) P-values are from parametric GSTs for profile panels; (**C-E, G-I, K-N**) P-values are from Mann–Whitney U tests for constituent biomarkers. Bars show medians with interquartile ranges. Sample sizes (control, PD): (**A, J-N**) 117, 163; (**B–E**) 175, 175; (**F-I**) 117, 165.

Collectively, the PD-status analyses therefore identified distributed but selective abnormalities across microbial function, epithelial injury/bacterial translocation, and myeloid activation rather than a global abnormality spanning all measured gut-related processes.

### PD-Spouse Sensitivity Analysis

The household-matched sensitivity analysis yielded a narrower pattern than the community-control comparison. Across available dyads (35-48 pairs depending on profile), neither the overall gut-related 24-biomarker profile (paired t-test: P=0.257; permutation P=0.258, **Table S5**) nor any broad parent mechanistic domain GST analyses reached nominal P<0.05: microbiome/dysbiosis (paired-t test: P=0.110), mucosa gut barrier/inflammation (P=0.350), and systemic inflammation/immune activation (P=0.835; **Table S5**). At, among the eight focused mechanistic clusters evaluated by GST, only the microbial functional-group profile differed significantly (paired t-test: P=0.030, Wilcoxon: P=0.016, and permutation: P=0.033; **Table S6**). This exploratory analysis was less informative for other profiles because paired sample sizes were small and variable. Nonetheless, the microbial functional-group signal remained detectable under household matching, whereas the other PD-control signals did not.

### PD Clinical Severity (State): Overall and Microbiome Profiles

The pattern across study-defined PD clinical states was different from the PD-control pattern. At the overall gut-related profile analysis, the 24-biomarker profile showed borderline and method-dependent evidence of variation across mild, moderate, and severe PD (GST P=0.053; Kruskal-Wallis P=0.036; permutation P=0.037; **Fig 4A**; **Table 2**). Among the broad mechanistic domains evaluated by GST, only the microbiome/dysbiosis profile differed significantly (GST P=0.023; permutation P=0.016; **Fig 4B**; **Table 2**); while no significant differences were found in both mucosa gut barrier/inflammation (GST P=0.076; permutation P=0.093; **Fig 5A**; **Table 2**) and systemic inflammation/immune activation domains (GST P=0.226; permutation P=0.261; **Fig 6A**; **Table 2**). Within the broad mechanistic microbiome/dysbiosis domain, alpha-diversity was the only focused profile that differed across clinical states (GST P=0.027; permutation P=0.031; **Fig 4C**; **Table 2**). The individual alpha-diversity measures showed the clearest difference for evenness (P=0.014), with nominal P=0.050 for Shannon and Simpson indices and no difference in observed features (P=0.135) (**Fig 4D-G**). In contrast to the PD-control comparison, neither microbial functional-groups (GST P=0.108; **Fig 4H**) nor microbial functional-ratios (GST P=0.141; **Fig 4M**) differed across clinical states (**Table 2**). Among the individual functional-group constituent biomarkers, only mucin-degrading genera differed nominally (P=0.024), whereas putative total SCFA-producing genera, putative butyrate-producing genera, and pathobiont genera did not (**Fig 4I-L**); neither derived functional-ratio differed individually (**Fig 4N-O**).

**Figure 4.**
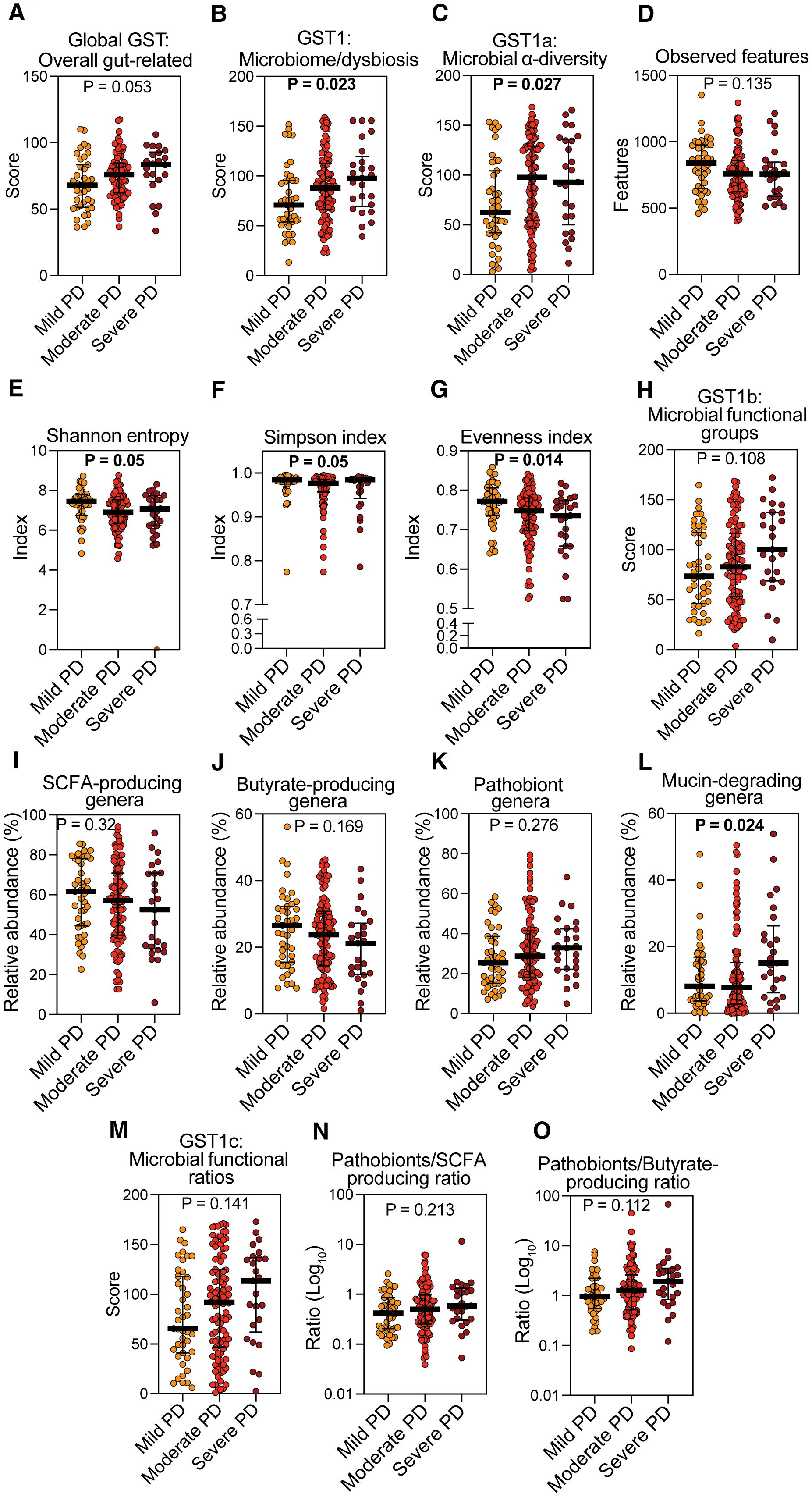
Overall and microbiome-related biomarker profiles across study-defined PD clinical states. (**A**) Overall gut-related 24-biomarker GST profile. (**B**) Broad mechanistic microbiome/dysbiosis profile. (**C-G**) Focused microbial alpha-diversity profile and constituent measures. (**H-L**) Focused microbial functional-group profile and constituent measures. (**M-O**) Derived microbial functional-ratio profile and constituent ratios. Clinical-state groups combine H&Y stage and medication status and should not be interpreted as pure disease-severity strata. Higher GST scores indicate greater prespecified abnormality after directional orientation and rank transformation. (**A-C, H, M**) P-values are from parametric GSTs for profile panels; (**D-G, I-L, N-O**) P-values are from Kruskal-Wallis tests for constituent biomarkers. Bars show medians with interquartile ranges. Sample sizes (mild, moderate, severe): (**A**) 39, 86, 22; (**B–O)** 43, 105, 25.

**Figure 5.**
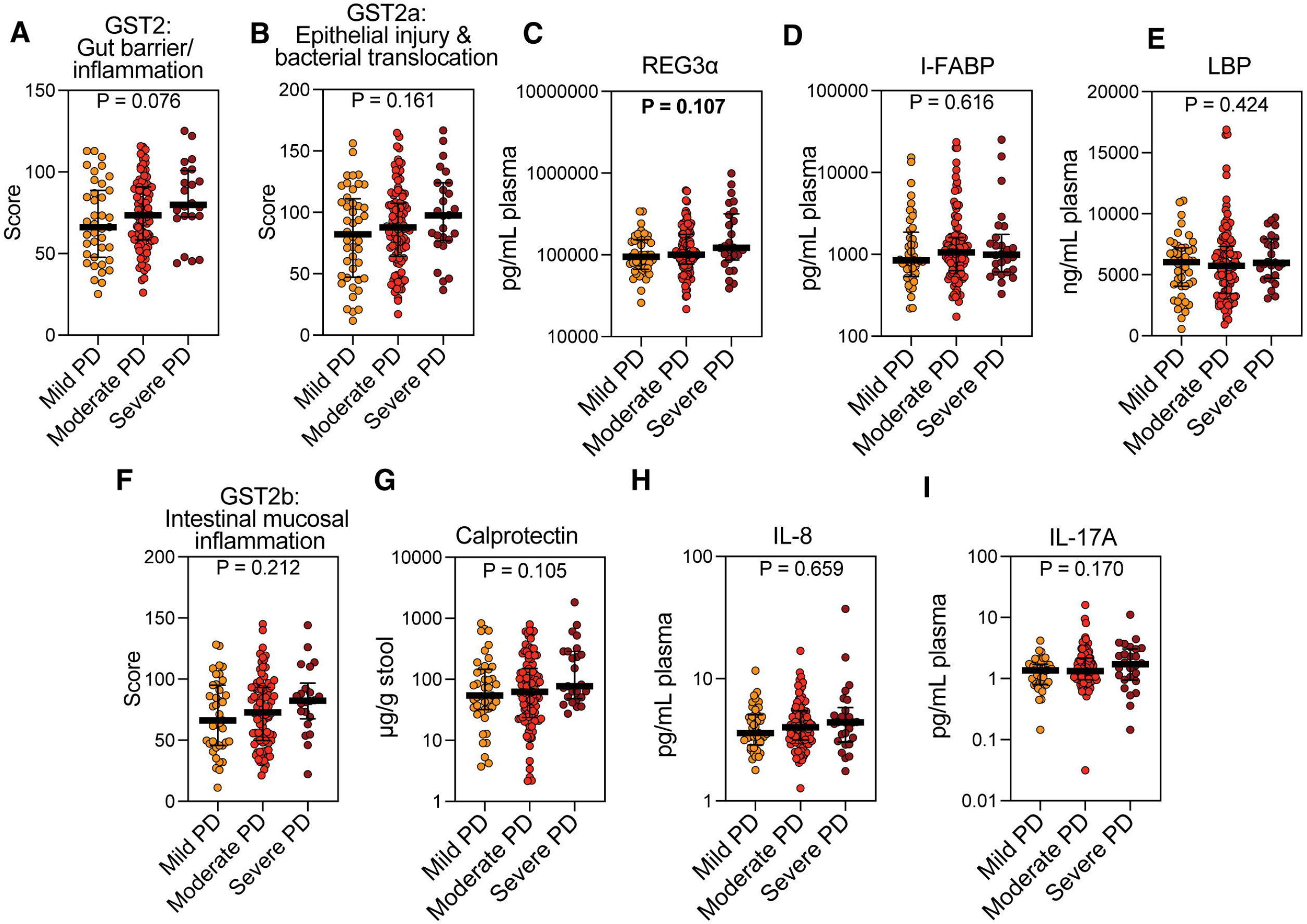
Mucosa gut leakage and inflammatory biomarker profiles across study-defined PD clinical states. (**A**) Broad mechanistic mucosa gut leakage/inflammation profile. (**B-E**) Focused epithelial injury/bacterial translocation profile and constituent biomarkers. (**F-I**) Focused intestinal mucosal-inflammation profile and constituent biomarkers. Clinical-state groups combine H&Y stage and medication status. (**A-B, F**) P-values are from parametric GSTs for profile panels; (**C-E, G-I**) P-values are from Kruskal-Wallis tests for constituent biomarkers. Bars show medians with interquartile ranges. Sample sizes (mild, moderate, severe): (**A, F-I**) 39, 86, 22; (**B–E)** 44, 105, 26.

**Figure 6.**
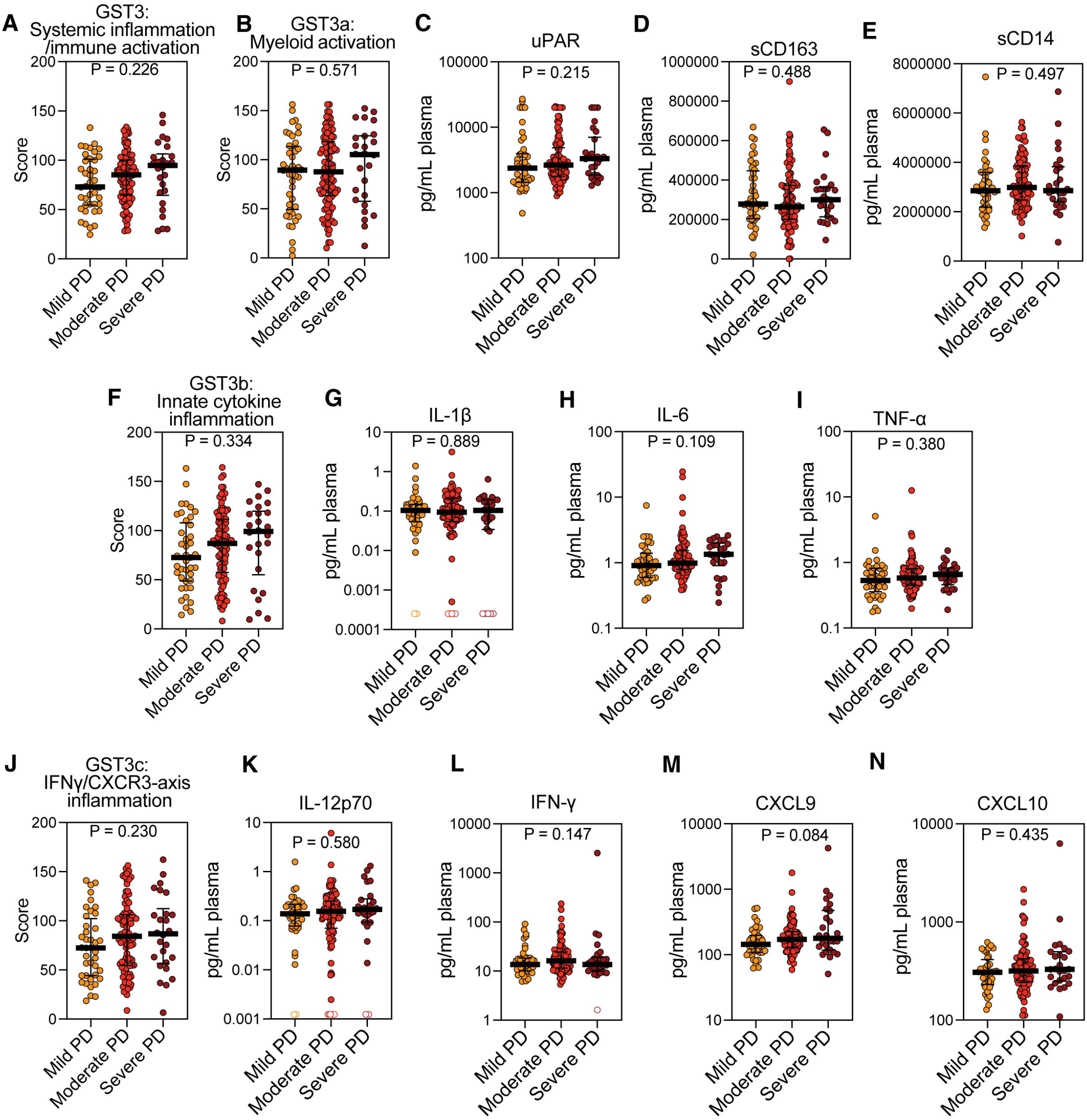
Systemic inflammatory and immune activation biomarker profiles across study-defined PD clinical states. (**A**) Broad mechanistic systemic inflammation/immune-activation profile. (**B-E**) Focused myeloid-activation profile and constituent biomarkers. (**F-I**) Focused innate-cytokine inflammation profile and constituent biomarkers. (**J-N**) Focused IFN-γ/CXCR3-axis inflammation profile and constituent biomarkers. Clinical-state groups combine H&Y stage and medication status. Values below the limit of detection ( LOD), for IL-1β, IL-12p70, and IFN-γ are colored coded in their respective figures. (**A-B, F, J**) P-values are from parametric GSTs for profile panels; (**C-E, G-I, K-N**) P-values are from Kruskal-Wallis tests for constituent biomarkers. Bars show medians with interquartile ranges. Sample sizes (mild, moderate, severe): (**A, J-N**) 40, 97, 26; (**B-E**) 44, 105, 26; (**F-I**) 41, 98, 26.

### PD Clinical Severity (State): Mucosa Gut Leakage/Inflammation and Systemic Inflammation/ Immune Activation Profiles

Neither mucosa gut leakage/inflammation or systemic inflammation/immune activation profiles showed evidence of a focused cluster difference across the study-defined PD severity groups-epithelial injury and bacterial translocation profile (GST P=0.161) and intestinal mucosal inflammation profile (GST P=0.212) were both null, and none of REG3A, I-FABP, LBP, stool calprotectin, IL-8, or IL-17A reached P<0.05 (**Fig 5B-I**; **Table 2**). Similarly, myeloid-activation (GST P=0.571), innate cytokine inflammation (GST P=0.334), and IFN-γ/CXCR3-axis inflammation (GST P=0.230) did not differ across PD clinical states, and none of their constituent plasma markers reached P<0.05 (**Fig 6B-N**; **Table 2**). This contrasted with the PD–control analysis, in which epithelial injury/bacterial translocation and myeloid activation emerged as prominent significant profiles.

### Secondary and Sensitivity Analyses of PD Clinical State

In age- and sex-adjusted models treating mild, moderate, and severe PD as ordered study-defined clinical states, the oriented alpha-diversity abnormality score increased with each one-level increase in clinical state (beta=0.280, 95% CI 0.043 to 0.517; P=0.021; **Table S3**). Because alpha-diversity measures were reverse-oriented before GST scoring, a higher score corresponds to lower raw diversity. Microbial functional-group and functional-ratio scores showed weaker trends (P=0.098 and P=0.100, respectively), while all mucosa gut leakage/inflammation and systemic inflammation/immune activation profiles had P>0.05 (**Table S3**). In the complete-case multivariate response model including all eight focused clusters, no individual domain reached P<0.05; the smallest P-value was 0.095 for epithelial injury and bacterial translocation (**Table S4**).

When disease severity was defined solely by H&Y stages 1-2 versus 3-5, neither overall gut-related biomarker profile GST analysis nor the broad mechanistic domains GST analysis reached P<0.05: overall gut-related GST P=0.159, microbiome/dysbiosis P=0.088, mucosa gut barrier/inflammation P=0.051, and systemic inflammation/immune activation P=0.321 (**Table S7**). At the focused cluster GST analysis, microbial functional-groups were borderline by the parametric GST (P=0.050) but not by permutation (P=0.053), and all other focused clusters were nonsignificant (**Table S8**).

For continuous MDS-UPDRS Part III, the age- and sex-adjusted separate models showed a nominal association with the broad microbiome/dysbiosis score (beta=1.805, 95% CI 0.036 to 3.574; P=0.046), but not with the overall gut-related profile (P=0.539), mucosa gut barrier/inflammation (P=0.230), or systemic inflammation/immune activation (P=0.125) (**Table S9**). No focused cluster scores reached P<0.05 in the separate models, and no focused cluster score reached P<0.05 in the joint model; epithelial injury/bacterial translocation showed the smallest joint-model P value (P=0.062) (**Table S9**).

### Exploratory Neurofilament light chain (NfL)-Burden Analysis

176 control and 176 PD-44 mild, 106 moderate, 26 severe had NFL measurement. For this exploratory analyses, PD participants were stratified using the 25th and 75th percentiles of the NfL distribution calculated exclusively within the PD cohort: low (≤91.1 pg/mL), intermediate (>91.1 to ≤180.46 pg/mL), and high (>180.46 pg/mL) (**Tables S10-11**). GST scores differed across cohort-defined low, intermediate, and high NfL groups for the GST overall gut-related profile analysis (GST P=0.003; permutation P<0.001), microbiome/dysbiosis (GST and permutation both P<0.001), and systemic inflammation/immune activation (GST P=0.027; permutation P=0.019), but not for mucosa gut leakage/ inflammation (GST P=0.235; **Table S10**).

At the focused cluster GST analysis, alpha-diversity differed (GST and permutation both P=0.002), microbial functional-groups showed GST P=0.050 and permutation P=0.040, and IFN-γ/CXCR3-axis inflammation showed GST P=0.052 and permutation P=0.033; other clusters did not reach nominal P<0.05 (**Table S11**). These exploratory tests were unadjusted for age and multiple comparisons; their association with NfL burden cannot be interpreted as evidence of progression or an independent association with neurodegeneration. Additionally, these cohort-derived thresholds were used for exploratory comparisons and do not represent clinically validated disease severity categories^42, 43^.

## DISCUSSION

This study was designed to access a large biological sample from PD patients and neurologically healthy controls to determine whether accessible gut-related biomarkers in PD form a global peripheral signature or instead reveal selective biological abnormalities. Further, we studied whether the gastrointestinal biomarker pattern associated with PD as a diagnosis is the same as the pattern associated with its clinical severity state. To both questions, the answer is a selective pattern of biomarker differences at both levels, but the diagnostic and severity patterns were distinctive with only partial overlap. PD status was not associated with a uniform shift across the 24-marker overall gut-related profile. Instead, microbiome/dysbiosis provided the clearest broad-domain signal, while focused epithelial injury/bacterial translocation and myeloid-activation focused clusters identified additional PD-associated abnormalities. Clinical state showed a different architecture: microbiome/dysbiosis again emerged at the broad parent level, but alpha-diversity rather than putative microbial functional composition was the clearest focused feature, and the gut barrier and myeloid profiles that distinguished PD from controls did not track the primary clinical-state severity groupings.

The PD-control results illustrate why a multi-level analytic design is biologically informative. A negative overall GST does not mean that overall gut-related biology is normal in PD. Rather, the abnormalities were distributed unevenly across the measured system. Within the microbiome, the signal was concentrated in putative microbial functional composition: PD participants had lower putative total anti-inflammatory SCFA- and butyrate-producing genera, higher mucin-degrading genera, and an altered pathobiont-to-butyrate-producing ratio, whereas alpha-diversity was not different between PD subjects and controls. Within the mucosa gut leakage/inflammation domain, whereas the full domain showed no difference between PD and controls, the specific epithelial injury/bacterial translocation profile was strongly different, driven most clearly by REG3A; stool calprotectin, both markers of gut inflammation, also differed even though the broader intestinal mucosal-inflammation profile did not. Likewise, the systemic immune domain analysis was globally null, yet the myeloid-activation profile differed markedly. These findings are consistent with selective biomarker associations of selective microbial, epithelial, and innate-immune perturbations rather than a single global “Parkinson gut” phenotype.

Importantly, the stronger broad-domain microbiome signal should not be interpreted as establishing biological primacy of the microbiome over the intestinal mucosa or immune system. Admittedly, the three parent domains were not interrogated at equivalent biological depth. Microbiome measures were obtained directly from stool microbial sequencing, whereas barrier and immune biology were represented mainly by circulating or fecal surrogate biomarkers rather than direct measurements of tissue structure, epithelial transcription, or functional permeability. The present study therefore has the advantage of comparing the signal carried by currently accessible biomarker panels, but it does not cover the full biological contribution in balanced format for each gut compartment.

The PD clinical state findings were related but distinct from the PD-trait findings. Microbiome/dysbiosis was the only broad domain that differed across the primary mild, moderate, and severe PD groups, but the focused signal shifted from putative microbial functional composition in the PD-control comparison to alpha-diversity measures across clinical states. The clinical-state model supported lower alpha-diversity indices with increasing state, whereas barrier and immune profiles remained nonsignificant. However, the primary state construct combines H&Y stage with medication status. PD medications can independently influence gut microbiome composition, making clinical severity phenotypes and treatment difficult to separate in this cross-sectional comparison.^4^ The H&Y-only analysis did not reproduce a statistically significant microbiome domain effect, and the MDS-UPDRS Part 3 score analysis showed only a nominal association with the broad microbiome/dysbiosis GST score. Accordingly, these analyses support PD study-defined clinical severity state-associated microbiome variation but do not establish a monotonic microbiome marker of PD progression.

The NfL analysis adds a distinct exploratory dimension. NfL is a blood biomarker of axonal neurodegeneration/neuroaxonal injury that has been associated with disease severity and future clinical progression in PD.^42, 43^ In this cohort, GST scores differed across cohort-defined NfL concentration groups not only with the overall 24 gut-related profile, but also with both microbiome/dysbiosis and systemic inflammation/immune activation profiles. At the focused mechanistic clusters assessment, alpha-diversity was the clearest result, with weaker or method-dependent evidence for microbial functional-groups and the IFN-γ/CXCR3 axis. This pattern raises the hypothesis that gut microbial and systemic immune abnormalities may covary with neurodegeneration burden, but the cross-sectional grouping, age dependence of NfL, unadjusted multiplicity, and absence of temporal data preclude inference about directionality or disease progression.

Stool and plasma can be collected repeatedly and in large sample size PD population, which makes these profiles suitable for prospective evaluation in settings where direct intestinal sampling is impractical. The present cross-sectional data does not establish diagnostic or prognostic performance and should not be interpreted as validated predictors. Instead, within a personalized, precision medicine framework, these findings identify candidate biomarker profiles for longitudinal evaluation in at-risk or prodromal cohorts for associations with PD onset, and in established PD for associations with progression, patient heterogeneity, or treatment response. The findings are also compatible with experimental and early clinical work showing that gut-directed interventions can alter microbiota, microbial metabolites, inflammatory readouts, and PD-relevant phenotypes, while not establishing that any single pathway identified here is a therapeutic target.^44, 45^

Several limitations are important. First, the cross-sectional design precludes conclusions about temporal sequence, prediction, or causality. Second, the study-defined clinical states combine H&Y stage and medication status and therefore are not pure disease-stage strata; the severe group and several sensitivity-analysis groups were relatively small. Third, PD and control groups differed in age, sex, and racial composition, while the primary GST comparisons were unadjusted for these covariates. These imbalances may correlate with diet, geography, socioeconomic exposures, medication use, and other factors relevant to microbiome and inflammatory measurements. Fourth, biomarker missingness produced different analytic samples across profiles, preventing formal decomposition of the broad tests into focused components. Fifth, P values were not adjusted for multiplicity, so specifically focused clusters GST analysis, focused individual-biomarker analyses, and sensitivity analyses should be regarded as exploratory. Sixth, the biological domains were measured at different depths, and circulating markers cannot fully characterize tissue-level mucosal biology. Seventh, the spouse analyses had smaller and variable paired samples. Eighth, taxonomically derived microbial functional groupings represent putative functions rather than direct measurements of microbial metabolic activity, and relative-abundance differences do not establish differences in absolute bacterial abundance. Finally, the study lacks an external replication cohort.

New avenues of future studies emerge from our study by the demonstration that accessible stool and plasma biomarkers can capture selective gut-related signatures of PD status and clinical state. Rather than focusing on a single global mechanistic abnormality, new studies for differentiating PD from controls, particularly as research focuses on prodromal or early diagnosis, can target microbiome-derived measures that have provided the most consistent broad-domain signal among the studied markers. Focused epithelial-injury/bacterial-translocation and myeloid-activation profiles identified herein can also be mechanistic PD Trait targets. For future studies on disease progression and clinical decline, alpha-diversity and microbiome-related markers and microbe-intestinal mucosa interaction can be potential candidates to follow. Exploratory NfL analyses further suggest that broader gut-related and systemic immune patterns may accompany neurodegeneration /neuroaxonal-injury burden.

## Supporting information

GTDB Taxonomic Classifier

## Acknowledgments

This work was supported in part by the National Institute of Diabetes and Digestive and Kidney Diseases, National Institutes of Health, under Award U01DK140923. Also, this research was supported in part by philanthropic funding from Mr. Larry and Mrs. Barbara Fields, Mr. Philip and Mrs. Helen Glass, and Mr. Harlan Berk. Rush is a designated Parkinson’s Foundation Clinical Center of Excellence.

## Author Contributions

A.K. conceived and designed the study, data interpretation, wrote the first draft and then edited the manuscript; SL performed GST data analysis, data interpretation; P.A.E. microbiota data analysis; statistical analysis, managed data recording, data interpretation; R.M.V interpreted the data, and edited the manuscript; M.S. laboratory measures, interpreted the data; AN microbiota data analysis, data interpretation; MV patient recruitment, data interpretation; APB laboratory measures, interpreted the data; LBG laboratory measures, interpreted the data; SJG microbiota data analysis, data interpretation; M.A.M. interpreted the data, and edited the manuscript; SL designed the GST analysis, interpreted the data, and edited the manuscript; C.G.G. designed the study, interpreted the data, and edited the manuscript. All authors reviewed and approved the final manuscript and have access to the data and the study design.

## Data Availability Statement

Reference the Source Data file. Microbiome data are available in the NCBI BioProject database under accession number PRJNA1511883. The classifier script used for data analysis is provided as a Supplementary Document. Additional data is available on request to corresponding author.

