## Supplemental Methods for "Selective Gut-Related Biomarker Profiles Define Parkinson Disease Trait and State"

**Supplementary Materials**

**Supplementary Methods**

*Blood Collection*

Plasma was previously collected by a RUMC phlebotomist into 10-mL K₂EDTA tubes (BD Vacutainer, #366643). Within 1-hour of collection, blood tubes were centrifuged at 800 x g for 15 minutes (acceleration setting 9; brake setting 9), plasma aliquoted and stored at −80°C.

*Stool Collection*

Stool specimens were previously collected by participants at home within 24–72 hours of their study visit at RUMC using the BD GasPak anaerobic system (Becton, Dickinson and Company; #260683). Participants recorded stool form, using the Bristol Stool Scale, scored as constipation (types 1–2), normal stool (types 3–4), or loose stool/diarrhea (types 5–7), and verified by the study clinical coordinator. Stool samples were stored at −80°C.

*Microbiome Profiling*

Genomic DNA was extracted from stool using the chemagic DNA Stool 200 Kit H96 (CMG-1076; Revvity) on a chemagic 360 automated extraction instrument according to the manufacturer's instructions. Samples underwent mechanical disruption using a TissueLyser II (Qiagen) at 30 Hz for two 5-minute cycles separated by a 2-minute rest period. Near-full-length bacterial 16S rRNA gene amplicons were generated using a previously described two-stage PCR protocol.^1^ The first PCR amplified the bacterial 16S rRNA gene using 27F and 1492R primers,^2^ followed by a second PCR incorporating PacBio Kinnex linkers and combinatorial indices.^3^ Amplified products were pooled, purified, and processed for Kinnex library preparation before sequencing on a PacBio Revio platform. Amplicon preparation was performed at the Genomics and Microbiome Core Facility (GMCF), while Kinnex library preparation and sequencing were conducted at the University of Illinois Urbana-Champaign (UIUC).

Near-full-length 16S rRNA gene sequences were processed using QIIME 2 (version 2025.7).^4^ Raw sequencing reads underwent initial quality assessment using SeqKit,^5^ followed by primer trimming with Cutadapt. Quality-filtered reads were denoised using DADA2 (q2-dada2-ccs plugin) to infer amplicon sequence variants (ASVs).^6^ Taxonomic classification was performed using a Naive Bayes classifier trained against the Genome Taxonomy Database (GTDB), release 226, using the majority dereplication method.^7, 8^ The classifier-generation script is provided in the Supplementary file. Potential reagent-derived contaminants were identified using the prevalence-based method implemented in the R package *decontam*,^9^ with default settings, by comparing ASV prevalence between negative controls and biological specimens. ASVs classified as potential contaminants were removed before downstream analyses. Alpha-diversity ASV counts were rarefied to 34,000 sequences per stool sample by random subsampling without replacement.
