## Supplementary figures and images for "Selective Gut-Related Biomarker Profiles Define Parkinson Disease Trait and State"

### Supplemental Figure 1

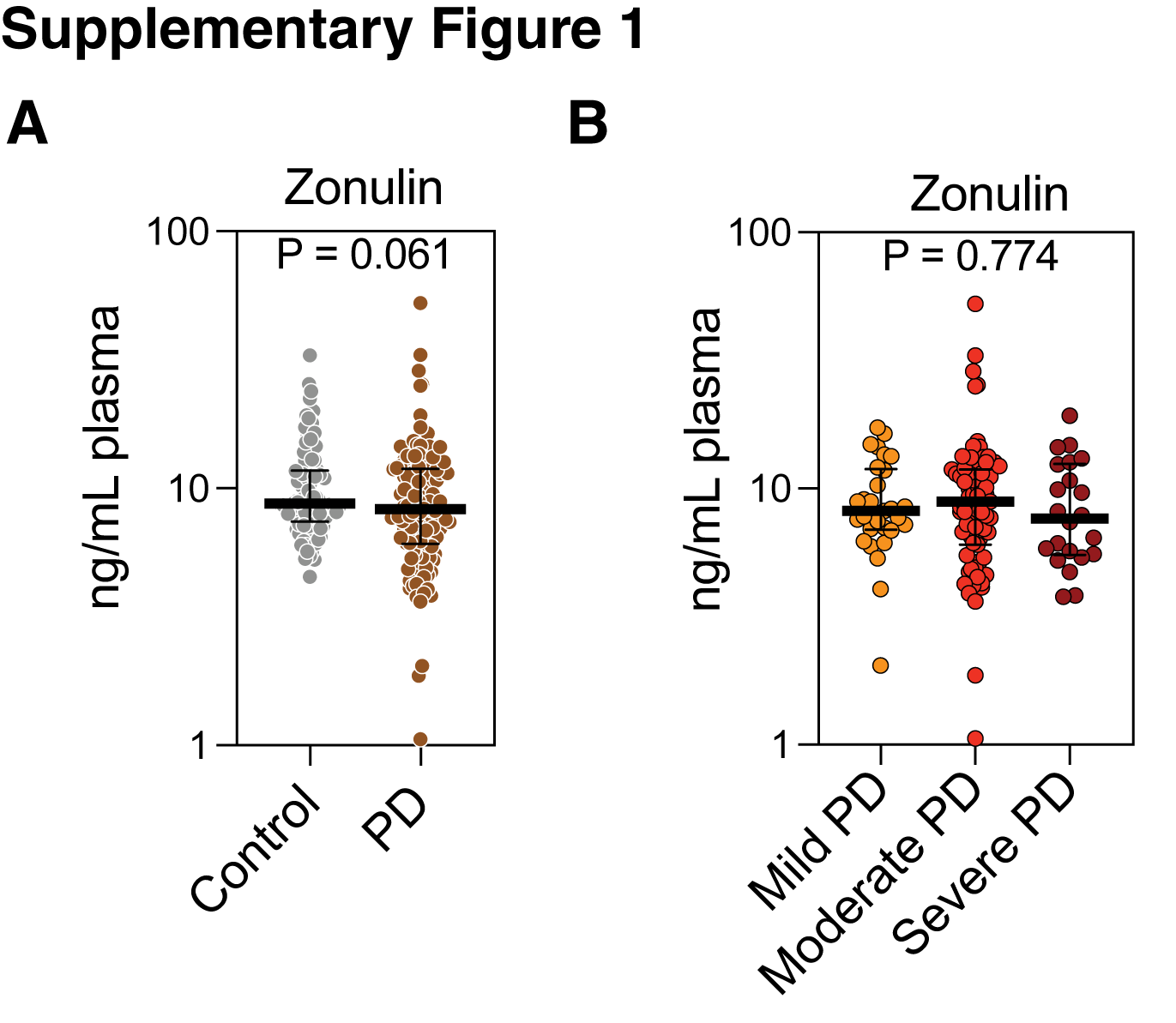
