## Supplemental Table 1 for "Selective Gut-Related Biomarker Profiles Define Parkinson Disease Trait and State"

**Supplementary Table S1. Demographics of study participants according to the paired PD-spouse.**

| **Variable** | **Spouse** | **Paired PD** | **Paired PD severity** | | |
| --- | --- | --- | --- | --- | --- |
|  | **Spouse (N=48)** | **Paired PD (N=48)** | **Mild (N=10)** | **Moderate (N=30)** | **Severe (N=8)** |
| Age | 62.5 [9.25] | 66 [9.75] | 66.5 [7.5] | 61.5 [11.75] | 68 [7.25] |
| Gender |  |  |  |  |  |
| Male | 15 (31.2%) | 34 (70.8%) | 8 (80.0%) | 19 (63.3%) | 7 (87.5%) |
| Female | 33 (68.8%) | 14 (29.2%) | 2 (20.0%) | 11 (36.7%) | 1 (12.5%) |
| Race |  |  |  |  |  |
| Caucasian | 45 (93.8%) | 45 (93.8%) | 8 (80.0%) | 29 (96.7%) | 8 (100.0%) |
| African American | 1 (2.1%) | 1 (2.1%) | 0 (0.0%) | 1 (3.3%) | 0 (0.0%) |
| Asian | 2 (4.2%) | 2 (4.2%) | 2 (20.0%) | 0 (0.0%) | 0 (0.0%) |
| Hispanic | 0 (0.0%) | 0 (0.0%) | 0 (0.0%) | 0 (0.0%) | 0 (0.0%) |
| Ethnicity |  |  |  |  |  |
| Non-Hispanic or Latino | 47 (97.9%) | 45 (93.8%) | 10 (100.0%) | 27 (90.0%) | 8 (100.0%) |
| Hispanic | 1 (2.1%) | 3 (6.2%) | 0 (0.0%) | 3 (10.0%) | 0 (0.0%) |
| Note: Age is summarized as median [IQR] in the table. Spouse: Age mean (SD) = 63.458 (8.262); Paired PD: Age mean (SD) = 65.062 (8.247); Mild: Age mean (SD) = 67.2 (4.78); Moderate: Age mean (SD) = 63.133 (8.815); Severe: Age mean (SD) = 69.625 (7.689). Categorical variables are summarized as n (%) on indented category rows. Percentages use group-specific denominators. BMI was excluded because of high missingness. | | | | | |
