## Supplemental Table 2 for "Selective Gut-Related Biomarker Profiles Define Parkinson Disease Trait and State"

**Supplementary Table S2. Biomarker composition and scoring direction of the gut biomarker GST profiles and corresponding gut-domain scores.**

| **Biological domain** | **GST subset** | **Score** | **Biological subset** | **Constituent biomarkers and scoring direction** |
| --- | --- | --- | --- | --- |
| Microbiome/dysbiosis | GST1a | D1 | Microbial alpha-diversity | Shannon index (lower); Simpson index (lower); observed features (lower); Pielou’s evenness (lower) |
| Microbiome/dysbiosis | GST1b | D2 | Microbial functional groups | Total SCFA-producing genera (lower); Total butyrate-producing genera (lower); Total pathobiont-producing genera (higher); Total mucin-degrading genera (higher) |
| Microbiome/dysbiosis | GST1c | D3 | Microbial functional ratios | Pathobiont/SCFA ratio (higher); Pathobiont/Butyrate ratio (higher) |
| Mucosa gut leakage/inflammation | GST2a | G1 | Epithelial injury and bacterial translocation | REG3A (higher); I-FABP (higher); LBP (higher) |
|  | GST2b | G2 | Intestinal mucosal inflammation | IL-8 (higher); IL-17A (higher); stool calprotectin (higher) |
| Systemic inflammation/immune activation | GST3a | S1 | Myeloid activation | uPAR (higher); sCD163 (higher); sCD14 (higher) |
| Systemic inflammation/immune activation | GST3b | S2 | Innate cytokine inflammation | IL-1β (higher); IL-6 (higher); TNF-α (higher) |
| Systemic inflammation/immune activation | GST3c | S3 | IFN-γ/CXCR3-axis inflammation | IL-12p70 (higher); IFN-γ (higher); CXCL9 (higher); CXCL10 (higher) |

Note: Biomarkers were oriented before rank aggregation so that higher oriented ranks represented greater hypothesized PD-associated abnormality. Variables listed as “lower” were reverse-oriented for GST scoring, whereas those listed as “higher” were retained in the adverse direction. Marker directions, including higher LBP as the adverse direction, were specified before the reported analyses and applied consistently across Levels 1-3. All Level 3 focused profiles are exploratory and were not adjusted for multiplicity. Abbreviations: CXCR3, C-X-C motif chemokine receptor 3; GST, Global Statistical Test; I-FABP, intestinal fatty acid-binding protein; IFN, interferon; IL, interleukin; LBP, lipopolysaccharide-binding protein; REG3A, regenerating islet-derived protein 3α; SCFA, short-chain fatty acids; sCD14, soluble CD14; sCD163, soluble CD163; TNF, tumor necrosis factor; uPAR, urokinase-type plasminogen activator receptor.
