## Supplemental Table 3 for "Selective Gut-Related Biomarker Profiles Define Parkinson Disease Trait and State"

**Supplementary Table S3. The associations of ordered PD clinical-state group with gut-domain scores.**

| **Biological domain** | **Score** | **n** | **β (95% CI)** | **P value** |
| --- | --- | --- | --- | --- |
| Microbiome/dysbiosis | D1 | 173 | 0.280 (0.043 to 0.517) | **0.021** |
|  | D2 | 173 | 0.200 (-0.037 to 0.436) | 0.098 |
|  | D3 | 173 | 0.198 (-0.038 to 0.434) | 0.100 |
| Mucosa gut leakage/inflammation | G1 | 175 | 0.184 (-0.059 to 0.427) | 0.137 |
|  | G2 | 147 | 0.145 (-0.108 to 0.397) | 0.260 |
| Systemic inflammation/immune activation | S1 | 175 | 0.074 (-0.156 to 0.305) | 0.526 |
|  | S2 | 165 | 0.153 (-0.094 to 0.400) | 0.224 |
|  | S3 | 163 | 0.103 (-0.146 to 0.352) | 0.416 |

Abbreviations: CI, confidence interval; H&Y, Hoehn and Yahr; PD, Parkinson disease. Each row is from a separate linear regression model with the listed standardized gut-domain score as the outcome and ordered study-defined PD clinical-state group (mild=0, moderate=1, severe=2) as the predictor of interest, adjusted for age and sex. β represents the estimated change in the standardized score associated with a one-level increase in the ordered study-defined clinical-state group. Because the ordering combines H&Y stage with medication status, this analysis is interpreted as an exploratory ordered clinical-state contrast rather than a pure disease-severity trend. D1-D3 corresponds to GST1a-GST1c; G1-G2 to GST2a-GST2b; and S1-S3 to GST3a-GST3c. This secondary clinical-state analysis is exploratory. All P-Values are two-sided and unadjusted for multiple comparisons.
