## Supplemental Table 4 for "Selective Gut-Related Biomarker Profiles Define Parkinson Disease Trait and State"

**Supplementary Table S4. The associations of ordered PD clinical-state group with gut-domain outcomes in a multivariate response model.**

| **Biological domain** | **Response** | **Gut-domain model** | |
| --- | --- | --- | --- |
|  |  | **β (95% CI)** | **P value** |
| Microbiome/dysbiosis | D1 | 0.205 (-0.047 to 0.457) | 0.112 |
|  | D2 | 0.103 (-0.149 to 0.355) | 0.425 |
|  | D3 | 0.106 (-0.148 to 0.359) | 0.415 |
| Mucosa gut leakage/inflammation | G1 | 0.212 (-0.035 to 0.459) | 0.095 |
|  | G2 | 0.145 (-0.106 to 0.395) | 0.260 |
| Systemic inflammation/immune activation | S1 | 0.032 (-0.211 to 0.276) | 0.796 |
|  | S2 | 0.100 (-0.163 to 0.363) | 0.457 |
|  | S3 | 0.091 (-0.167 to 0.349) | 0.492 |

Abbreviations: CI, confidence interval; H&Y, Hoehn and Yahr; PD, Parkinson disease. The multivariate linear regression model jointly included D1-D3, G1-G2, and S1-S3 as eight standardized responses, adjusted for age and sex, and included 147 participants. β represents the estimated change in the corresponding response associated with a one-level increase in the ordered study-defined PD clinical-state group (mild=0, moderate=1, severe=2). Because this ordering combines H&Y stage with medication status, it is interpreted as an exploratory ordered clinical-state contrast rather than a pure disease-severity trend. Residual covariance was estimated jointly within the model. This secondary clinical-state analysis is exploratory. All P-Values are two-sided and unadjusted for multiple comparisons.
