## Supplemental Table 5 for "Selective Gut-Related Biomarker Profiles Define Parkinson Disease Trait and State"

**Supplementary Table S5. Level 1 overall and Level 2 intermediate GST results in paired PD-spouse sensitivity analyses.**

| **Level** | **Analytic profile** | **No. markers** | **Paired n** | **Paired t** | **df** | **t P** | **Wilcoxon** | **Asym. P** | **Perm. P** |
| --- | --- | --- | --- | --- | --- | --- | --- | --- | --- |
| Level 1 | Overall gut-related | 24 | 35 | 1.15 | 34 | 0.257 | 380.00 | 0.291 | 0.258 |
| Level 2 | Microbiome/dysbiosis | 8 | 39 | 1.64 | 38 | 0.110 | 465.50 | 0.087 | 0.114 |
|  | Mucosa gut leakage/inflammation | 6 | 35 | 0.95 | 34 | 0.350 | 335.00 | 0.527 | 0.344 |
|  | Systemic inflammation/immune activation | 10 | 36 | -0.21 | 35 | 0.835 | 313.00 | 0.759 | 0.828 |

Abbreviations: GST, Global Statistical Test; PD, Parkinson disease. Level 1 is the overall 24-biomarker gut-related profile. Level 2 uses the non-overlapping intermediate sets described in the main text: microbiome/dysbiosis (8 unique biomarkers from D1 and D2), mucosa gut leakage/inflammation (6 unique biomarkers from G1-G2), and systemic inflammation/immune activation (10 unique biomarkers from S1-S3). Paired t tests, Wilcoxon signed-rank tests, and two-sided sign-flip permutation tests were used. The number of paired dyads varies with biomarker availability; the “No. markers” column is not participant sample size. This paired sensitivity analysis is exploratory. All P-Values are two-sided and unadjusted for multiple comparisons.
