## Supplemental Table 6 for "Selective Gut-Related Biomarker Profiles Define Parkinson Disease Trait and State"

**Supplementary Table S6. Level 3 focused paired GST profiles between participants with PD and spousal controls.**

| **Biological domain** | **GST profile** | **Paired dyads, n** | **Paired tests** | | |
| --- | --- | --- | --- | --- | --- |
|  |  |  | **Paired t P** | **Wilcoxon P** | **Permutation P** |
| Microbiome/dysbiosis | GST1a: Microbial alpha-diversity | 39 | 0.797 | 0.961 | 0.790 |
|  | GST1b: Microbial functional groups | 39 | **0.030** | **0.016** | **0.033** |
|  | GST1c: Microbial functional ratios | 39 | 0.103 | 0.101 | 0.106 |
| Mucosa gut leakage/inflammation | GST2a: Epithelial injury and bacterial translocation | 48 | 0.990 | 0.850 | 0.989 |
|  | GST2b: Intestinal mucosal inflammation | 35 | 0.848 | 0.838 | 0.851 |
| Systemic inflammation/immune activation | GST3a: Myeloid activation | 48 | 0.236 | 0.338 | 0.236 |
|  | GST3b: Innate cytokine inflammation | 37 | 0.421 | 0.305 | 0.424 |
|  | GST3c: IFN-γ/CXCR3-axis inflammation | 36 | 0.953 | 1.000 | 0.952 |

Abbreviations: GST, Global Statistical Test; PD, Parkinson disease. Difference direction was PD minus spouse. Wilcoxon results are from paired signed-rank tests; permutation P-values are from two-sided sign-flip tests on paired differences. All paired focused-profile analyses are sensitivity analyses and exploratory; P-values are unadjusted for multiplicity. All P-values are two-sided.
