## Supplemental Table 7 for "Selective Gut-Related Biomarker Profiles Define Parkinson Disease Trait and State"

**Supplementary Table S7. Level 1 and Level 2 gut-domain GST profiles comparing H&Y stages 1-2 versus stages 3-5 among participants with Parkinson disease.**

| **Level** | **Analytic profile** | **No. markers** | **Sample n (n1,n2)** | **GST F** | **df** | **GST P** | **K-W H** | **Asym. P** | **Perm. P** |
| --- | --- | --- | --- | --- | --- | --- | --- | --- | --- |
| Level 1 | Overall gut-related | 24 | 147(125,22) | 2.01 | 1, 145 | 0.159 | 3.11 | 0.078 | 0.078 |
| Level 2 | Microbiome/dysbiosis | 8 | 173(148,25) | 2.94 | 1, 171 | 0.088 | 2.81 | 0.093 | 0.089 |
|  | Mucosa gut leakage/inflammation | 6 | 147(125,22) | 3.87 | 1, 145 | 0.051 | 3.41 | 0.065 | 0.066 |
|  | Systemic inflammation/immune activation | 10 | 163(137,26) | 0.99 | 1, 161 | 0.321 | 1.15 | 0.283 | 0.286 |
| Abbreviations: df, degrees of freedom; GST, Global Statistical Test; H&Y, Hoehn and Yahr; PD, Parkinson disease. | | | | | | | | | |
| H&Y groups were defined among participants with PD as stages 1-2 versus stages 3-5. Level 1 is the overall 24-biomarker gut-related profile. Level 2 uses the non-overlapping broad sets described in the main text. | | | | | | | | | |
| The sample size column is shown as n(n1,n2), where n is the total analytic sample size, n1 is the H&Y stage 1-2 group size, and n2 is the H&Y stage 3-5 group size. | | | | | | | | | |
| For the two-group comparison, the GST statistic is the parametric group-comparison statistic, and the nonparametric statistic is Kruskal-Wallis H; permutation P-Values are based on the same rank-based group comparison. Analytic sample sizes vary across profiles because of biomarker availability. This H&Y severity analysis is exploratory. All P-Values are two-sided and unadjusted for multiple comparisons. | | | | | | | | | |
