## Supplemental Table 8 for "Selective Gut-Related Biomarker Profiles Define Parkinson Disease Trait and State"

**Supplementary Table S8. Level 3 focused gut-domain GST profiles comparing H&Y stages 1-2 versus stages 3-5 among participants with Parkinson disease.**

|  |  |  | **GST results** | | | |
| --- | --- | --- | --- | --- | --- | --- |
| **Biological domain** | **GST profile** | **Sample size** | **GST F** | **P value** | **Kruskal-Wallis H** | **Permutation P** |
| Microbiome/dysbiosis | GST1a: Microbial alpha-diversity | 173(148,25) | 0.66 | 0.419 | 0.62 | 0.432 |
|  | GST1b: Microbial functional groups | 173(148,25) | 3.89 | **0.050** | 3.69 | 0.053 |
|  | GST1c: Microbial functional ratios | 173(148,25) | 2.38 | 0.125 | 2.18 | 0.135 |
| Mucosa gut leakage/inflammation | GST2a: Epithelial injury and bacterial translocation | 175(149,26) | 2.60 | 0.108 | 2.15 | 0.146 |
|  | GST2b: Intestinal mucosal inflammation | 147(125,22) | 2.67 | 0.104 | 2.28 | 0.135 |
| Systemic inflammation/immune activation | GST3a: Myeloid activation | 175(149,26) | 0.79 | 0.377 | 0.77 | 0.390 |
|  | GST3b: Innate cytokine inflammation | 165(139,26) | 0.69 | 0.408 | 1.32 | 0.259 |
|  | GST3c: IFN-γ/CXCR3-axis inflammation | 163(137,26) | 0.52 | 0.470 | 0.55 | 0.470 |
| Abbreviations: GST, Global Statistical Test; H&Y, Hoehn and Yahr; PD, Parkinson disease. | | | | | | |
| H&Y groups were defined among participants with PD as stages 1-2 versus stages 3-5. Profiles contain the focused gut-domain biomarkers. Kruskal-Wallis statistics are shown with permutation P-Values as robustness analyses. | | | | | | |
| The sample size column is shown as n(n1,n2), where n is the total analytic sample size, n1 is the H&Y stage 1-2 group size, and n2 is the H&Y stage 3-5 group size. | | | | | | |
| All Level 3 focused-profile H&Y analyses are exploratory and were not adjusted for multiplicity. Analytic sample sizes varied by profile because of biomarker availability. All P-Values are two-sided. | | | | | | |
