## Supplemental Table 9 for "Selective Gut-Related Biomarker Profiles Define Parkinson Disease Trait and State"

**Supplementary Table S9. Associations of derived gut-domain scores with MDS-UPDRS Part III score.**

|  |  |  | **Separate model** | | **Joint model** | |
| --- | --- | --- | --- | --- | --- | --- |
| **Biological domain** | **Score** | **n** | **Separate model Beta (95% CI)** | **Separate model P value** | **Joint model Beta (95% CI)** | **Joint model P value** |
| Overall gut-related |  | 144 | 0.604 (-1.336 to 2.544) | 0.539 |  |  |
| Microbiome/dysbiosis |  | 170 | 1.805 (0.036 to 3.574) | **0.046** |  |  |
| Mucosa gut leakage/inflammation |  | 144 | 1.141 (-0.732 to 3.015) | 0.230 |  |  |
| Systemic inflammation/immune activation |  | 156 | -1.407 (-3.206 to 0.393) | 0.125 |  |  |
| Microbiome/dysbiosis | D1 | 170 | 1.154 (-0.600 to 2.908) | 0.196 | 0.366 (-1.612 to 2.344) | 0.715 |
|  | D2 | 170 | 1.688 (-0.064 to 3.441) | 0.059 | 1.324 (-0.684 to 3.331) | 0.194 |
|  | D3 | 170 | 1.569 (-0.186 to 3.325) | 0.079 |  |  |
| Mucosa gut leakage/inflammation | G1 | 168 | 1.530 (-0.179 to 3.238) | 0.079 | 2.123 (-0.112 to 4.357) | 0.062 |
|  | G2 | 144 | 0.407 (-1.550 to 2.364) | 0.681 | 1.556 (-0.827 to 3.938) | 0.199 |
| Systemic inflammation/immune activation | S1 | 168 | -0.414 (-2.254 to 1.426) | 0.658 | -0.979 (-3.241 to 1.284) | 0.394 |
|  | S2 | 158 | -1.415 (-3.225 to 0.394) | 0.124 | -1.468 (-3.873 to 0.936) | 0.229 |
|  | S3 | 156 | -1.278 (-3.085 to 0.529) | 0.164 | -1.630 (-3.849 to 0.589) | 0.149 |
| Abbreviations: CI, confidence interval; MDS-UPDRS, Movement Disorder Society-Unified Parkinson’s Disease Rating Scale; PD, Parkinson disease. | | | | | | |

Separate models were multiple linear regression models assessing the association between continuous MDS-UPDRS Part III item sum and each standardized gut-related score (Level 1, Level 2, and Level 3), adjusted for age and sex. The joint model included D1, D2, G1-G2, and S1-S3 as predictors, together with age and sex. D3 was excluded from the joint model because it is constructed from measures used in D2. Beta represents the estimated change in MDS-UPDRS Part III score associated with a one-standard-deviation increase in the corresponding standardized gut-related score. This secondary motor-severity analysis is exploratory. All P-Values are two-sided and unadjusted for multiple comparisons.
