## Supplemental Table 10 for "Selective Gut-Related Biomarker Profiles Define Parkinson Disease Trait and State"

**Supplementary Table S10. Level 1 and Level 2 gut-domain GST profiles comparing low, intermediate, and high NfL-burden groups among participants with Parkinson disease.**

| **Level** | **Analytic profile** | **No. markers** | **Analytic n (low, intermediate, high)** | **GST F** | **df** | **GST P** | **K–W H** | **Asym. P** | **Perm. P** |
| --- | --- | --- | --- | --- | --- | --- | --- | --- | --- |
| Level 1 | Overall gut-related | 24 | 147 (39, 73, 35) | 6.04 | 2, 144 | **0.003** | 13.62 | 0.001 | **<0.001** |
| Level 2 | Microbiome/dysbiosis | 8 | 151 (40, 74, 37) | 7.65 | 2, 148 | **<0.001** | 14.47 | <0.001 | **<0.001** |
| Level 2 | Mucosa gut leakage/inflammation | 6 | 147 (39, 73, 35) | 1.46 | 2, 144 | 0.235 | 3.28 | 0.194 | 0.189 |
| Level 2 | Systemic inflammation/immune activation | 10 | 163 (41, 83, 39) | 3.71 | 2, 160 | **0.027** | 8.01 | 0.018 | **0.019** |

Abbreviations: GST, Global Statistical Test; NfL, neurofilament light chain; PD, Parkinson disease. NfL groups were defined within PD by the 25th and 75th percentile cut points: low, ≤91.1 pg/mL; intermediate, >91.1 to ≤180.46 pg/mL; high, >180.46 pg/mL. The group-specific analytic sample sizes vary across profiles. GST F is the parametric one-way ANOVA statistic; Kruskal–Wallis H and its permutation P-Value are robustness analyses. All comparisons are exploratory, unadjusted for age and for multiple testing.
