## Supplemental Table 11 for "Selective Gut-Related Biomarker Profiles Define Parkinson Disease Trait and State"

Supplementary Table S11: Level 3 focused gut-domain GST profiles comparing low, intermediate, and high NfL-burden groups among participants with Parkinson's disease.

|  |  |  | **GST results** | | | |
| --- | --- | --- | --- | --- | --- | --- |
| **Biological domain** | **GST profile** | **Sample size** | **GST F** | **P value** | **Kruskal-Wallis H** | **Permutation P** |
| Microbiome/dysbiosis | GST1a: Microbial alpha-diversity | 151(40,74,37) | 6.27 | **0.002** | 11.80 | **0.002** |
|  | GST1b: Microbial functional groups | 151(40,74,37) | 3.06 | **0.050** | 6.43 | **0.040** |
|  | GST1c: Microbial functional ratios | 151(40,74,37) | 2.42 | 0.093 | 4.68 | 0.097 |
| Mucosa gut leakage/inflammation | GST2a: Epithelial injury and bacterial translocation | 175(44,87,44) | 1.55 | 0.215 | 3.02 | 0.225 |
|  | GST2b: Intestinal mucosal inflammation | 147(39,73,35) | 0.72 | 0.491 | 1.37 | 0.503 |
| Systemic inflammation/immune activation | GST3a: Myeloid activation | 175(44,87,44) | 1.54 | 0.218 | 3.17 | 0.208 |
|  | GST3b: Innate cytokine inflammation | 165(42,83,40) | 1.90 | 0.153 | 3.94 | 0.138 |
|  | GST3c: IFN-γ/CXCR3-axis inflammation | 163(41,83,39) | 3.00 | 0.052 | 6.80 | **0.033** |
